# DIA-based quantitative proteomics reveal the protective effects of quercetin against atopic dermatitis via attenuating inflammation and modulating immune response

**DOI:** 10.64898/2026.01.08.698465

**Authors:** Meng Lu, Derong Chen, Xiongshun Liang, Liru Feng, Xiaoqian Liu, Xuqiao Hu, Wenxu Hong

## Abstract

Atopic dermatitis is a common inflammatory skin disorder characterised by recurrent eczematous lesions and intense itch. Quercetin is a naturally occurring flavonoid, exhibits antioxidant and anti-inflammatory properties. Previous studies have indicated its beneficial role in managing atopic dermatitis (AD); however, the precise underlying mechanism remains unclear. This study aimed to investigate the protective effects of quercetin against AD and to elucidate its potential mechanisms. We demonstrated that quercetin effectively suppressed the overexpression of pro-inflammatory effector factors—including interleukin (IL)-1β, thymic stromal lymphopoietin (TSLP) and chemokine (C-X-C motif) ligand 1/10/11 (CXCL1/10/11)—in immortalized human keratinocytes (HaCaT) stimulated by TNF-α/IFN-γ. Proteomic analysis revealed 88 differentially expressed proteins, suggesting that quercetin’s therapeutic action involves multiple pathways. Notably, the NOD-like receptor signaling pathway was identified as a key factor. The key proteins related to this pathway including inhibitor of nuclear factor kappa B kinase epsilon (IKBKE), indoleamine 2,3-dioxygenase 1 (IDO1) and chemokine (C-X-C motif) ligand 9 (CXCL9) showed a significant decrease in expression levels after quercetin treatment. Collectively, these results demonstrate that quercetin alleviates AD symptoms by attenuating inflammatory responses, likely through inhibition of the NOD-like receptor signaling pathway.

## 1. Introduction

Atopic dermatitis (AD) is a common chronic relapsing inflammatory disease with multifactorial etiology, characterized by epidermal barrier deficiency and immune overactivation, and dominated by Th2-related cytokine responses^1,2^. Global Burden of Disease studies have shown prevalence rates of 15% to 20% in children and up to 10% in adults, making atopic dermatitis the 15th most common nonfatal disease and the dermatologic condition with the highest burden of disease^3^. Glucocorticoids, calcineurin inhibitors and biopharmaceuticals have some problems such as side effects, poor efficacy in some patients and high cost^4–7^. Thus, natural compounds, which inhibit AD may prove to be promising agents for prostate cancer therapy^8^.

Quercetin is a naturally occurring flavonoid with antioxidant, anti-tumor, and anti-inflammatory activities. Previous studies have reported the effects of quercetin on alleviating AD symptoms^9–11^. Xing-hua Gao group’s results show that quercetin treatment can significantly improve the skin lesions of AD mice^11^. Nathaniel S Hwang’s research is consistent with this^9^. However, the target and mechanism of quercetin remain unclear due to lack of in-depth research and validation.

Proteomics has been widely used for identifying new molecular targets and elucidating the mechanisms of natural products. By comparing changes in protein expressions upon drug treatment, proteomics provides rich information on understanding mechanism-of-action of a drug and its toxicity^12^.

In order to enhance the understanding of the molecular mechanisms of quercetin in the treatment of AD, in this study, we investigated the effects of quercetin on the proteomic profile of human keratinocyte HaCaT cells. We showed that a negative regulator of inhibitor of nuclear factor kappa B kinase Subunit epsilon (IKBKE) and indoleamine 2,3-Dioxygenase 1 (IDO1), are two of the key regulators related to quercetin treatment. Our findings may aid improvement of translational application of quercetin and development of novel anti-AD drugs.

## 2 Materials and Methods

### Reagents

Quercetin (purity ≥98%) was provided by MedChemexpress Biotechnology (Shanghai, China), and it’s structure is shown in Figure 1A. Recombinant human TNF-α and recombinant human IFN-γ were purchased from Peprotech (London, UK). Cell Counting Kit-8 (CCK-8) was purchased from Beyotime (Shanghai, China).

**Figure 1.**
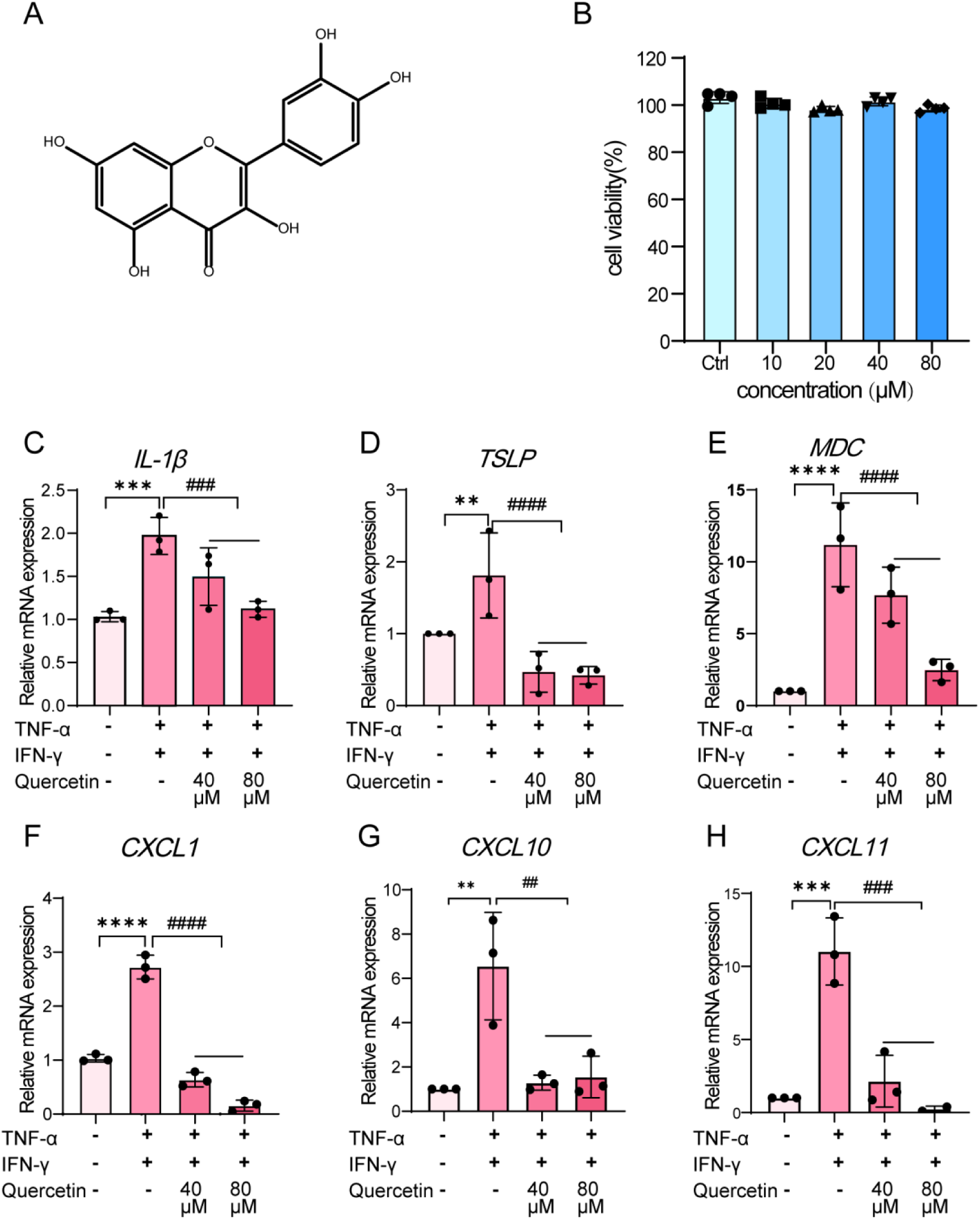
Effect of quercetin on cell viability and TNF-α/IFN-γ induced inflammation response in HaCaT. (A) The chemical structure of quercetin. (B) HaCaT cells were treated with different concentrations of quercetin(0, 10, 20, 40 and 80 μM) for 24 h, and cell viability was determined by CCK8 assay. (C-H) HaCaT cells were pretreated with 40 μM quercetin or 80 μM quercetin for 30 min and then exposed to 10 ng/mL TNF-α and 10 ng/mL IFN-γ for a further 24 h. Then mRNA expression of IL-1β, MDC, TSLP, MDC, CXCL1,CXCL10 and CXCL11 were detected by real-time qPCR. \*\**p* < 0.01, \*\*\**p* < 0.001, \*\*\*\**p* < 0.0001, ###*p* <0.001, ####*p* < 0.0001.

Antibodies against IL-1α, IL-1β, RANTES, IDO1, CXCL9, IKBKE, ICAM-1 and GAPDH were purchased from Proteintech (Wuhan, China). Total RNA Isolation Kit was obtained from Omega Bio-tek (Norcross, Georgia, USA). PrimeScript™ RT reagent Kit and TB Green® Premix Ex Taq^TM^ Ⅱ were purchased from Takara Bio (Tokyo, Japan). Bicinchoninic acid assay (BCA) kit was purchased from Thermo (Shanghai, China).

### Cell culture

Human keratinocyte HaCaT cells, purchased from Procell (Wuhan, China), were cultured in DMEM (Gibco, C11995500BT) containing 10 % fetal bovine serum (Gibco, A5669701) with 1% penicillin and streptomycin (Gibco, 15140-122) in a 5% CO2 incubator at 37°C.

HaCaT cells (5 × 10^5^ cells/well) in 6-well plates were treated with quercetin (40 and 80 μM) for 30 minutes, then the cells were co-treated with IFN-γ and TNF-α (all 10 ng/ml) in the presence or absence of quercetin for 24 h.

### Cell viability

HaCaT cells were plated at a density of 1 × 10^4^ cells/well in 96-well plates and treated with quercetin of gradient concentration and cultured 24 h in complete DMEM media. CCK-8 assay was used to detect the cell viability according to manufacturer’s protocol. CCK8 solution was added to each well and incubated for 4 h in incubator. And then, the optical density (OD) at 450 nm was detected through a full-wavelength multifunctional microplate reader (Thermo Scientific™ Varioskan™ LUX).

### Quantitative Real-time PCR (qRT-PCR)

Total RNA was extracted from cells using a total RNA extraction kit. Then use the kit reverse transcription to RNA into cDNA. qRT-PCR was performed as previously described. Gene expression was normalized to GAPDH mRNA. The primers used in this study are shown in Table 1.

**Table 1.**
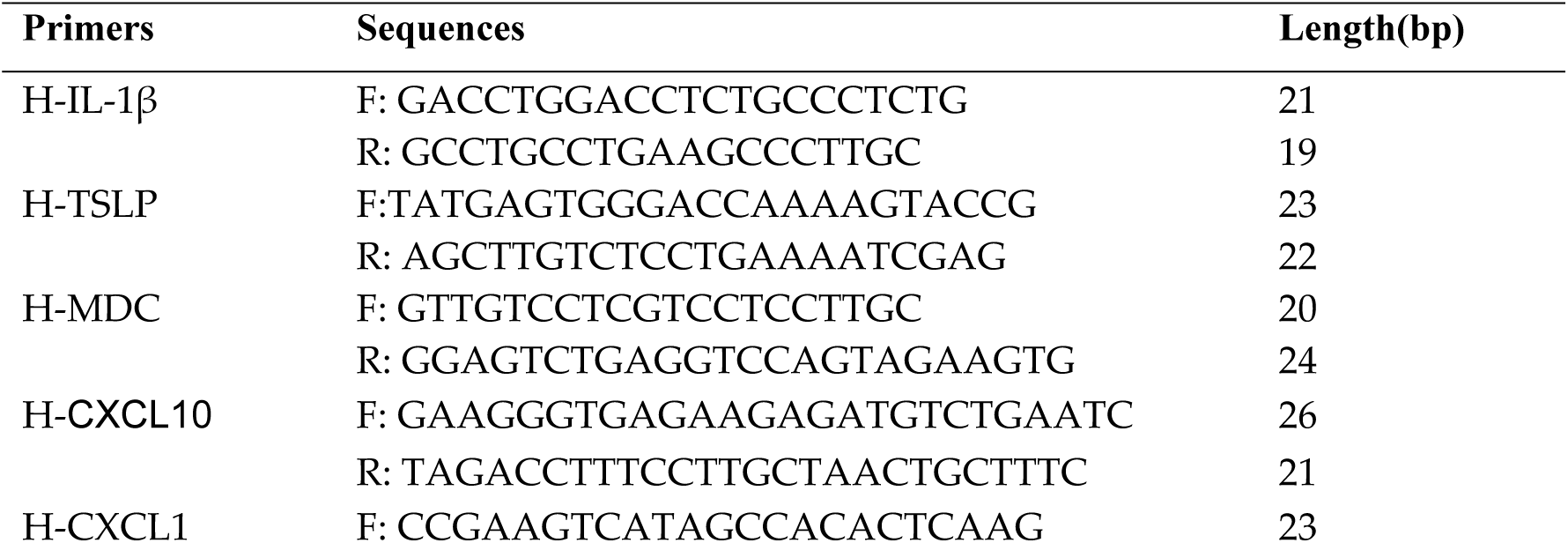
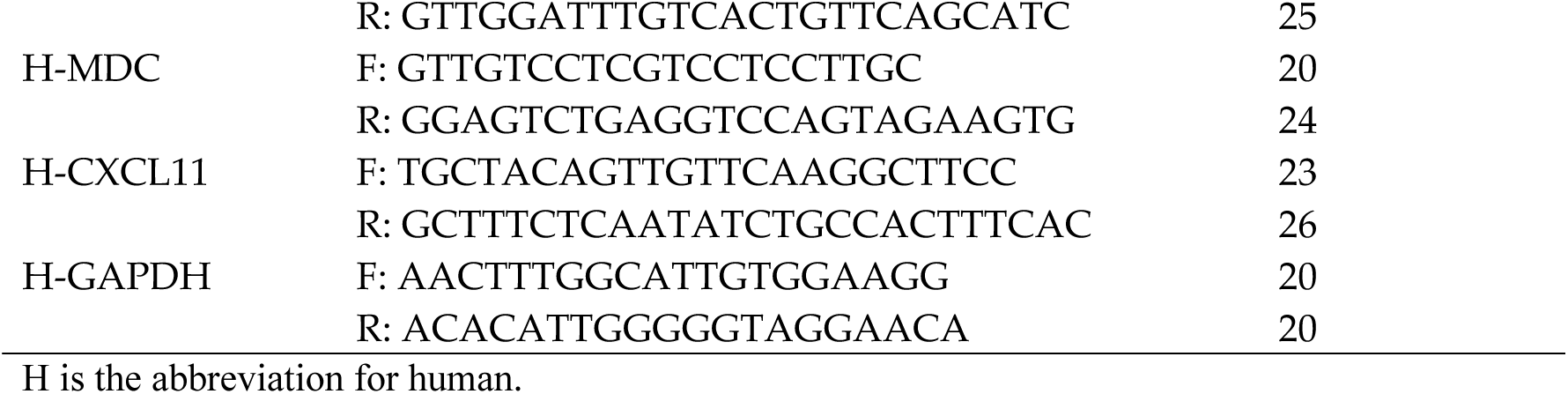
Sequences of primers used for quantitative real-time PCR (qRT-PCR).

### Western blot

The samples were isolated by 10% sodium dodecyl sulfate-polyacrylamide gel electrophoresis (SDS-PAGE) and transferred onto polyvinylidene fluoride membranes. The membranes were blocked with 5% nonfat milk containing Tris-buffered saline with 0.1% Tween 20 (TBST) for 1 h at RT, washed with TBST five times, and incubated with anti-IL-1α, anti-IL-1β, anti-RANTES, anti-IDO1, anti-CXCL9, anti-IKBKE, ICAM-1 and anti-GAPDH (diluted to 1:3000, Abcam) antibodies overnight. Following this, the membranes were washed again and incubated with the corresponding secondary antibodies for 1 h. The membrane was analyzed using enhanced chemiluminescence reagents, and the density of the bands was quantified using Image J.

### Protein sample extraction and pre-treatment

As previously described. Total protein was extracted using 8 M urea and The protein concentrations were determined with the BCA kit. Next, the extracted proteins were denatured, reduced, acylated, and digested. Briefly, protein samples were denatured in 8 M urea/25 mmol/L NH4HCO3, reduced in 10 mmol/L dithiothreitol (DTT), acylated in 50 mmol/L iodoacetamide (IAA), and added according to the mass ratio of trypsin: protein=1:40 Trypsin was added at a mass ratio of trypsin:protein=1:40, and the peptide samples were incubated at a constant temperature of 37°C in a water bath for 16 h. The peptide samples were recovered and spun dry under vacuum. The peptides were solubilized with 0.1% formic acid (FA)/water (H_2_O), and the concentration was detected by NanoVue and diluted to 500 ng/μL.

### LC-MS/MS setup for DIA

LC-MS/MS analysis was performed using an Easy-nLC 1200 liquid chromatograph and an Orbitrap Fusion mass spectrometer, with 0.1% FA/H2O as mobile phase A and 80% acetonitrile (ACN)/0.1% FA as mobile phase B. The flow rate of the pump was 2 μL/min, and the samples were loaded on to the analytical column Acclaim™ PepMap™ 100 C18 HPLC (75 μmx25cm, 3 μm) as the analytical column. PepMap™ 100 C18 HPLC (75 μmx25 cm, 3 μm) was used as the analytical column, and the separation flow rate was 0.3 μL/min. The peptides were eluted according to the following program: 0.1 ∼102 min: 5% B∼25% B; 102∼110 min: 25% B∼38% B; 110∼112 min: 38% B∼90% B; and finally 112 ∼120 min: 90% B. The peptides were extracted from the peptides at a flow rate of 2 μL/min. In addition, the spray voltage was 2.3 kV, the scanning range (m/z) was 300∼1500, the MS1 resolution was 60,000, and the maximum injection time was 50 ms; the MS/MS resolution was 15,000, and the maximum injection time was 100 ms.

### Label-free DIA quantification analysis

The mass spectrometry data were searched by DIA-NN, and the search conditions were set as follows: species: human (UniProt homo sapiens database); protease species: trypsin; maximum missed cleavage sites: 2; error range of primary parent ions: 10 ppm; error range of secondary fragment ions: 20 ppm; fixed modifications: cysteine ureido-methylation; variable modifications: methionine oxidation, protein N-terminal acetylation; False Discovery Rates (FDR) ≤ 05%. : methionine oxidation, protein N-terminal acetylation; False Discovery Rates (FDR) ≤ 0.05. The plausible proteomic data were log2 transformed using Label-free quantitation (LFQ) intensity data to fill in the missing values according to the normal distribution, and a two-tailed t-test was performed to separate each protein between the two groups of samples. A two-tailed t-test was performed between the samples and the corresponding p-value was calculated, with p < 0.05 as the significance indicator. In this study, proteins with a multiplicity of difference greater than 1.5 and p < 0.05, or with a multiplicity of difference less than 0.67 and p < 0.05, were regarded as differential proteins.

### Pathway and functional enrichment analyses

For biological process and pathway enrichment analysis, KEGG and Gene Ontology (GO) analysis was performed by metascape. Parameter settings: Min Overlap: 3, P Value Cutoff: 0.01, Min Enrichment: 1.5.

### Statistical analysis

Data was presented as the mean ± SEM, with (*) and (#) as notes. Student’s t-test for two-group comparisons and one-way ANOVA were used, followed by post-hoc tests for multiple comparisons among more than two groups. Differences were considered statistically significant at p < 0.05. GraphPad Prism (Version 8.0) was used for the statistical analysis.

## 3 Results

### 3.1 Effect of quercetin on cell viability and TNF-α/IFN-γ induced inflammation response in HaCaT

To evaluate the effect of quercetin on cell viability, HaCaT cells were treated with quercetin at concentrations of 0, 10, 20, 40, and 80 μM for 24 h, and cell viability was assessed using the CCK-8 assay. As shown in Figure 1B, quercetin did not exhibit detectable cytotoxicity at concentrations up to 80 μM. Based on these results, 40 and 80 μM were selected for subsequent experiments.

To assess the effect of quercetin on TNF-α/IFN-γ-induced inflammatory responses, HaCaT cells were pretreated with quercetin (40 or 80 μM) for 30 min and subsequently stimulated with TNF-α (10 ng/mL) and IFN-γ (10 ng/mL) for 12 h. The mRNA expression levels of inflammatory cytokines and chemokines were examined by quantitative real-time PCR. TNF-α/IFN-γ stimulation significantly increased the mRNA expression of IL-1β, TSLP, MDC, CXCL1, CXCL10, and CXCL11 compared with the control group. Quercetin pretreatment markedly reduced the expression of these inflammatory mediators in a concentration-dependent manner (Figure 1C–H).

### 3.2 General characteristics of the proteome of TNF-α/IFN-γ stimulated HaCaT cells

To further explore the potential mechanism of quercetin improving AD, we performed quantitative proteomic analysis on HaCaT cells in control (Ctrl), TNF-α/IFN-γ and Que-80 μM groups in three independent replicates for each group. We screened these differentially expressed proteins (DEPs) using the criteria of folding change multiple (FC) ≥ 1.5 or ≤ 0.67 and *p* < 0.05.

Proteome analysis showed that 1379 DEPs were identified in TNF-α/IFN-γ group and Ctrl group, including 391 up-regulated proteins and 988 down-regulated proteins.In addition, 190 DEPs were found between the Que-80 μM group and the TNF-α/IFN-γ group, including 78 up-regulated proteins and 112 down-regulated proteins (Figure 2). Furthermore, 88 DEPs were reversed in the Que-80 μM group compared with the TNF-α/IFN-γ group (Figure 2D).

**Figure 2.**
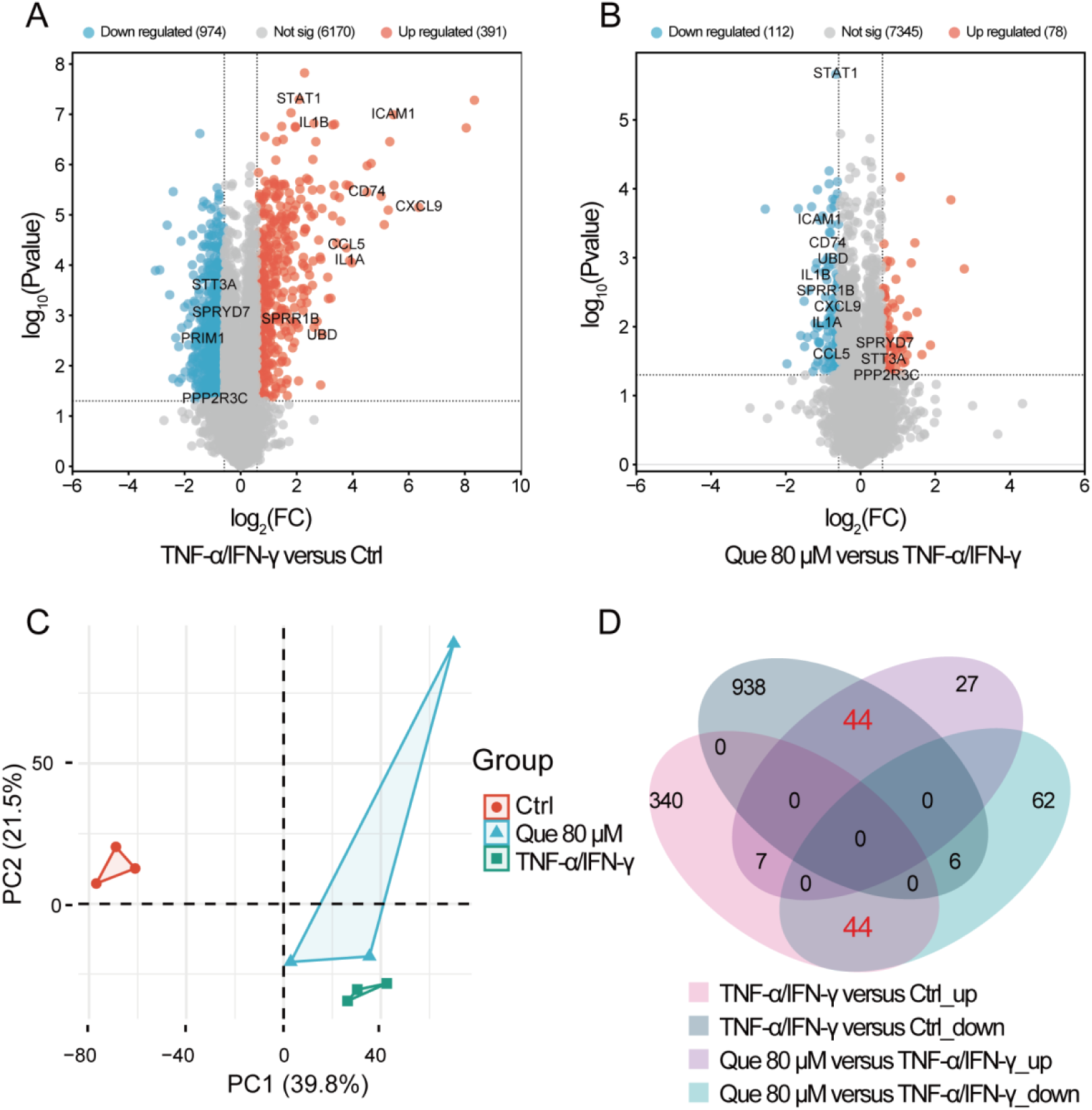
Lable-free proteomic analysis of control group, model group and quercetin treated group. (A) Volcano plots of DEPs in control and model groups. (B) Volcano plots of DEPs in model and quercetin-treated groups. (C) Principal component analysis (PCA). (D) Venn diagrams of up-regulated proteins and down-regulated proteins in model group compared to control versus up-regulated proteins and down-regulated proteins in quercetin-treated group compared to model.

Principal component analysis (PCA) of HaCaT cell proteome showed a clear separation between the Ctrl group and TNF-α/IFN-γ group. Proteins in the Que-80 μM group were located between the model group and the control group, suggesting a potential protective effect of Que-80 (Figure 2C).

### 3.3 Bioinformatics analysis of DEPs

Hierarchical clustering analysis demonstrated that the proteomic profiles of the quercetin-treated group were more similar to those of the control group than to the model group (Figure 3B). To further characterize the biological functions of the reversed proteins, Gene Ontology (GO) enrichment analysis was performed, revealing significant enrichment in cytokine-mediated signaling and inflammatory responses (Figure 3C). KEGG pathway analysis indicated that these proteins were mainly involved in NOD-like receptor and Toll-like receptor signaling pathways (Figure 3A).

**Figure 3.**
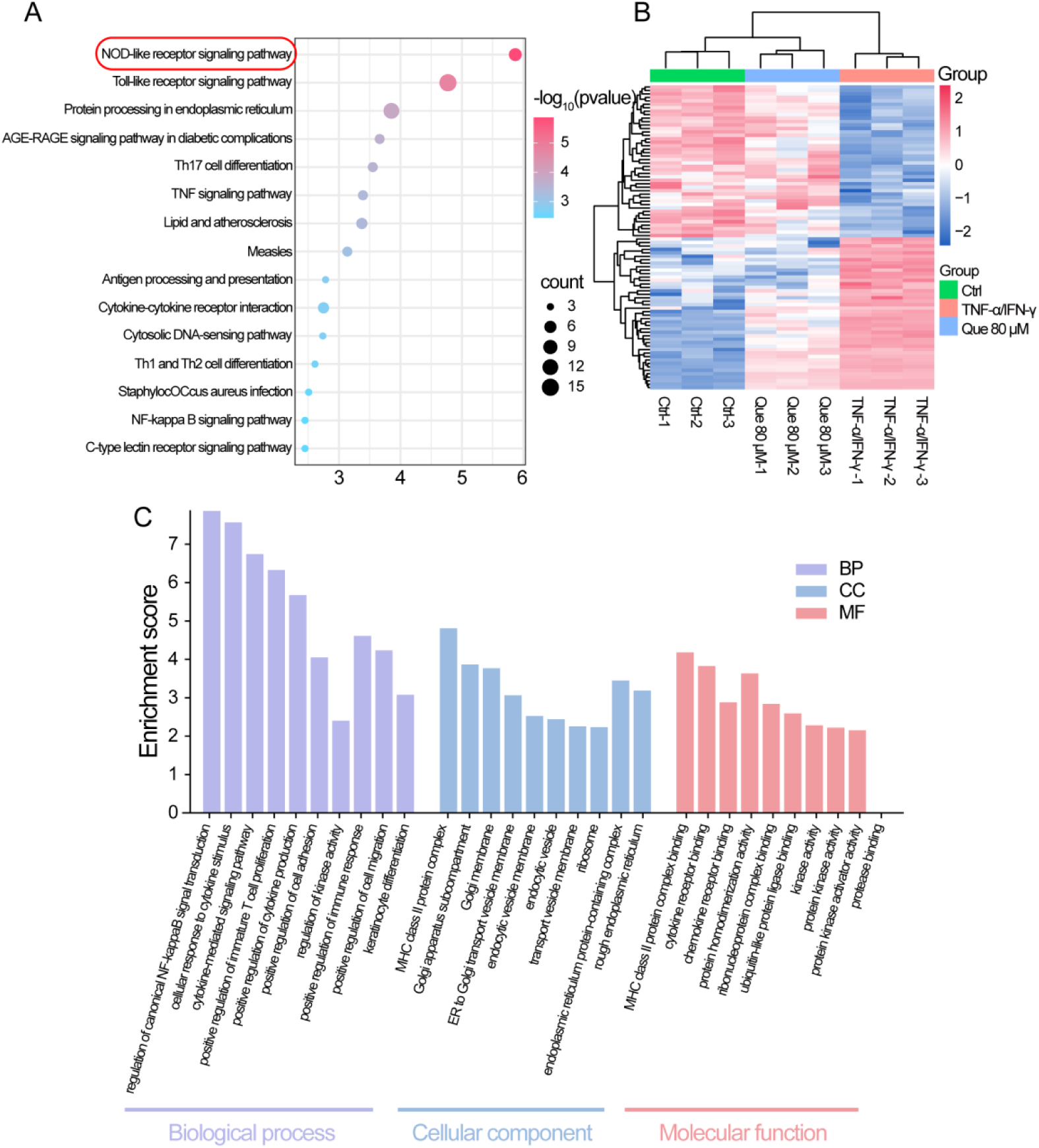
Bioinformatics analysis of DEPs. (A) KEGG pathway analysis of the 88 DEPs, (B) Heatmap, and (C) GO analysis of the 88 DEPs.

### 3.4 Validation of proteomic results by western blot

The protein expression levels of IKBKE, IDO1, CXCL9, and RANTES were significantly increased in TNF-α/IFN-γ-stimulated HaCaT cells and were markedly reduced by quercetin pretreatment, consistent with the proteomic analysis results (Figure 4).

**Figure 4.**
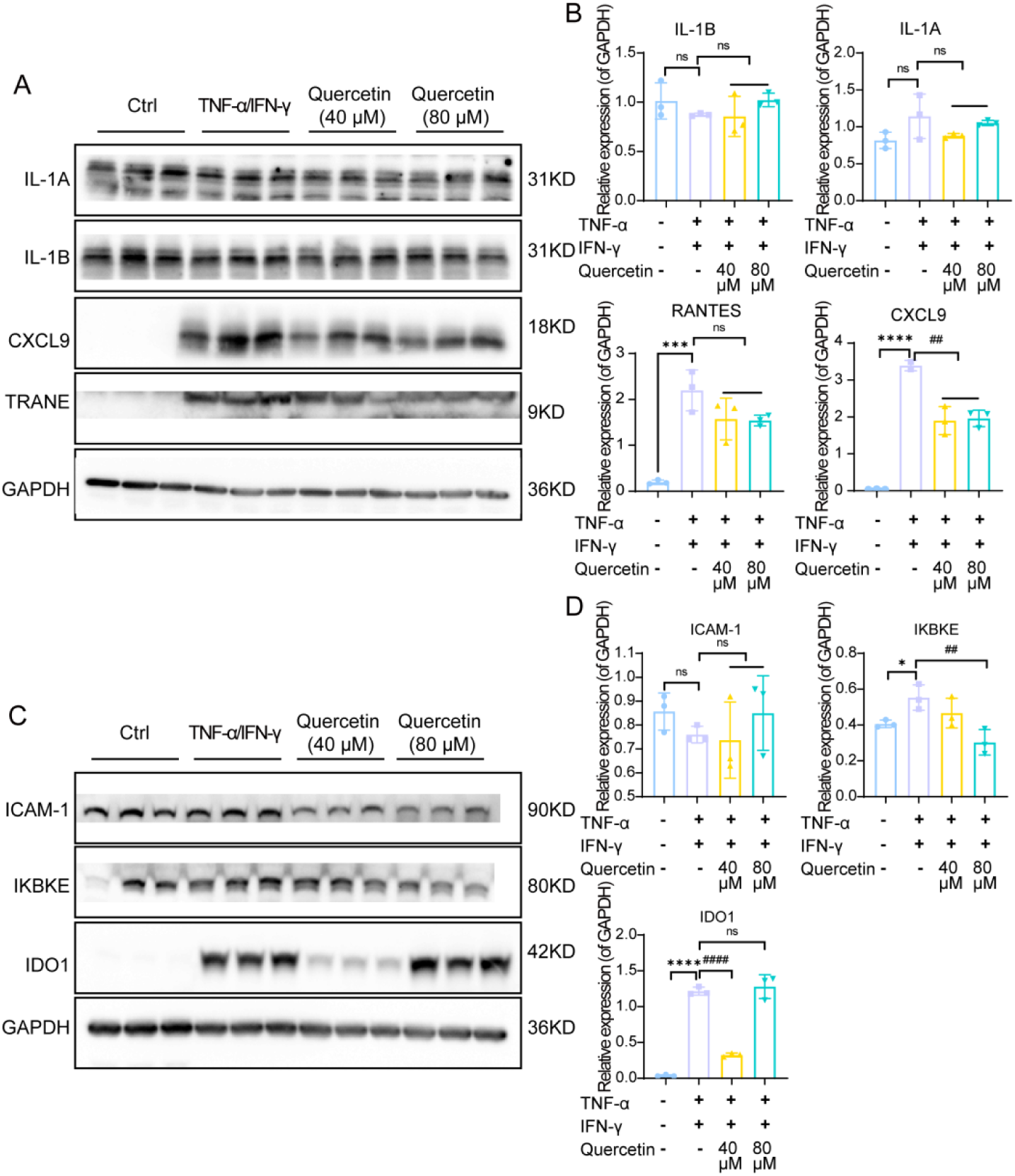
WB verified the key differential expression proteins. HaCaT cells were pretreated with quercetin (40 or 80 μM) for 30 min and then stimulated with TNF-α/IFN-γ for 24 h. Protein expression levels of IKBKE, IDO1, CXCL9, and RANTES were detected by western blotting and quantified relative to GAPDH.

## 4. Discussion

Keratinocytes are the main cell type in the epidermis, serving as both an external structural barrier and an active immune mediator for host defense against pathogens^13^. Studies confirm that Keratinocytes are atopic dermatitis in enhanced cells of the immune response, and skin immune response is the main derivatives and targets. When stimulated by external factors, epidermal keratinocytes release various pro-inflammatory cytokines and chemokines, further exacerbating skin inflammation^14^. During the occurrence and development of AD, activated keratinocytes produce various chemokines including CCL-type chemokines and CXCL-type chemokines. These chemokines recruit immune cells such as T cells and monocytes into the inflamed skin, further exacerbating skin inflammation^15–18^. Additionally, thymic stromal lymphopoietin (TSLP) secreted by keratinocytes can promote the differentiation of initial CD4^+^ T cells into Th2 cells, enabling them to produce allergic cytokines IL-4, IL-5, IL-13, and TNF-α^19,20^. In this study, we found that quercetin can inhibit the upregulation of mRNA expression of inflammatory cytokines IL-1β and chemokines TSLP, MDC, and CXCL1/10/11 in TNF-α/IFN-γ-stimulated HaCaT cells. These results indicate that quercetin has anti-inflammatory and immunomodulatory pharmacological properties, demonstrating its potential as a therapeutic drug for AD.

To further explore the potential mechanism of quercetin in the treatment of AD, we used quantitative proteomics study based on data independent acquisition. Through this study, we identified 1,379 proteins with defined quality and quantifiable properties. Among them, 88 proteins were found to be reversed by quercetin from the upregulation or downregulation caused by TNF-α/IFN-γ. These differentially expressed proteins (DEPs) served as the basis for our subsequent bioinformatics analysis. The KEGG pathway enrichment analysis revealed that the differentially expressed proteins were significantly enriched in the NOD-like receptor signaling pathway and the Toll-like receptor signaling pathway, indicating their crucial role in the therapeutic effect of quercetin on AD. Further identification of key proteins related to the NOD-like receptor signaling pathway and the Toll-like receptor signaling pathway, including CXCL9, IKBKE, IDO1, and RANTES (CCL5), revealed that the NLRP3 inflammasome plays a key role in the acute and chronic stages of AD, regulating the immune bias^13^ between Th2 and Th1. Additionally, NOD can collaborate with TLR and its related activation signaling pathways to initiate a potent antibacterial immune response^21^. These functions are crucial for maintaining immune homeostasis and defending against infections.

According to the report, nucleotide and oligomerization domain structure like receptor (NLRs), including five family, namely the NLRA, NLRB, NLRC, NLRP and NLRX, Plays an important role in a variety of autoimmune diseases^22^. In NLR family members, NLRP3 cohesion protein apoptosis related mottled protein (ASC) and the activity of enzyme original pro - caspase 1 form ternary protein complex inflammatory corpuscle, Inductive effect protein caspase 1 will pro - IL - 1 beta, and pro - IL - 18 cut into proinflammatory cytokines IL - 1 beta and IL - 18, causing inflammation and Th2, Th17 related immune imbalance^21^. Studies have found that the nuclear factor-κB (NF-κB) signaling pathway is crucial for the activation of basophils, eosinophils, and dermal fibroblasts in NOD2/TLR2 ligand-mediated AD-related inflammation^23,24^. NOD2/TLR2 activates the NF-κB signaling pathway, thereby accelerating the expression of pro-inflammatory cytokines and chemokines. Additionally, the IKBKE gene (also known as IKKε) encodes IKBKE, which is a key kinase of the IKK family. Research reports that when cells sense external pathogens or inflammatory signals, IKBKE is activated, subsequently phosphorylating IκB protein. This process leads to ubiquitination of IκB protein and eventual degradation by the proteasome, thereby releasing NF-κB transcription factors. The released NF-κB enters the cell nucleus and initiates the transcription of a series of genes related to immune responses and inflammation^25,26^.

Indoleamine 2,3-dioxygenase 1 (IDO1) is a rate-limiting enzyme with immunomodulatory properties, which degrades tryptophan (TRP) through the kynurenine pathway (KP). There is literature indicating that IDO1 can activate two main pathways: (1) By consuming TRP, it activates the general control non-repressive kinase 2 (GCN2K) pathway and inhibits the mammalian target of rapamycin (mTOR) signaling pathway, thereby leading to T-cell autophagy, anergy, and apoptosis; (2) The kynurenine (KYN) produced and its metabolites can activate the aryl hydrocarbon receptor (AhR), which helps maintain persistent immune tolerance by influencing the balance of Th1/Th2 and Th17/Tregs systems^27^. This suggests that controlling the expression of IDO1 may inhibit the development process of AD.

In conclusion, our findings indicate that quercetin significantly improves the inflammatory response in hacat cells induced by TNF-α/IFN-γ. The therapeutic effect of quercetin seems to be related to its ability to inhibit the expression of proteins such as IKBKE, IDO1, and CXCL9, which are involved in the NOD-like receptor signaling pathway. This suggests that these proteins have the potential to become effective targets for the treatment of AD. Our study has confirmed the anti-inflammatory property of quercetin in AD, laying the foundation for its clinical application.

### Author Contributions

Meng Lu: Writing-review & editing, Writing-original draft, Visualization, Methodology, Formal analysis, Data curation. Derong Chen: Writing-original draft, Visualization, Validation, Investigation, Data curation. Xiongshun Liang: Data curation, Investigation. Liru Feng: Data curation, Investigation. Xiaoqian Liu: Visualization, Validation, Investigation. Xuqiao Hu: Supervision, Project administration, Funding acquisition, Conceptualization. Wenxu Hong: Writing-review & editing, Supervision, Project administration, Funding acquisition.

## Acknowledgments

This work was supported by the Guangdong Province Medical Science and Technology Research Fund Project (A2024128).

